# Keywords to success: a practical guide to maximise the visibility and impact of academic papers

**DOI:** 10.1101/2023.10.02.559861

**Authors:** Patrice Pottier, Malgorzata Lagisz, Samantha Burke, Szymon M. Drobniak, Philip A. Downing, Erin L. Macartney, April Robin Martinig, Ayumi Mizuno, Kyle Morrison, Pietro Pollo, Lorenzo Ricolfi, Jesse Tam, Coralie Williams, Yefeng Yang, Shinichi Nakagawa

## Abstract

In a growing digital landscape, enhancing the discoverability and resonance of scientific articles is essential. Here, we offer ten recommendations to amplify the discoverability of studies in scientific databases. Particularly, we argue that the strategic use and placement of key terms in the title, abstract, and keyword sections can boost indexing and appeal. By surveying 237 journals in ecology and evolutionary biology, we found that current author guidelines may unintentionally limit article discoverability. Our survey of 5842 studies revealed that authors frequently exhaust abstract word limits — particularly those capped under 250 words. This suggests that current guidelines may be overly restrictive and not optimised to increase the dissemination and discoverability of digital publications. Additionally, 91.9% of studies used redundant keywords in the title or abstract, undermining optimal indexing in databases. We encourage adopting structured abstracts to maximise the incorporation of key terms in titles, abstracts, and keywords. In addition, we encourage the relaxation of abstract and keyword limitations in journals with strict guidelines, and the inclusion of multilingual abstracts to broaden global accessibility. These evidence-based recommendations to editors are designed to improve article engagement and facilitate evidence synthesis, thereby aligning scientific publishing with the modern needs of academic research.

## Introduction

Scientific articles serve as the primary method for disseminating research findings. Between 1980 and 2012, global scientific output was estimated to increase by 8 – 9% every year, implying a doubling of scientific evidence approximately every nine years [1]. Amid this burgeoning landscape, standing out becomes a research agenda in its own right. Ensuring that articles are well written and indexed in databases such as Scopus, or Web of Science is akin to laying the first bricks in the foundation — it is important for discoverability, but not sufficient. Many articles, despite being indexed, remain undiscovered. We argue that carefully crafting titles, abstracts, and keywords is a critical step to increase the visibility and impact of scientific research.

Titles, abstracts, and keywords are the primary marketing components of any scientific paper, and valorising these elements is crucial [2,3]. However, studies with appealing abstracts will not necessarily be discovered because of a lack of search engine optimisation. Search engine optimisation is the process of enhancing the findability of content by search engines. While often not discussed in the academic sphere, it is particularly relevant for scientific articles. To discover articles, academics use a combination of key terms in scientific literature databases or search engines, and most databases leverage algorithms to scan the words in titles, abstracts, and keywords to find matches. Failure to incorporate appropriate terminology could thus undermine readership. Search engines such as Google Scholar may look through the articles in their entirety. However, the same underlying principle remains - the absence of critical key terms means these articles would not surface in the search results. Keywords also play an important role in the search ranking process.

Choosing the right terms can often mean the difference between a study appearing at the top of the search results or being buried beneath a virtual pile of other documents. This is particularly important for databases that sort results by relevance, where the strategic use of keywords can significantly enhance the article’s visibility. Additionally, not including relevant keywords impedes a study’s inclusion in literature reviews and meta-analyses, which often rely on database searches based on key terms in titles, abstracts, and keywords [4,5].

Enhancing study discoverability is, however, ineffective if the abstract fails to engage the reader. Readers typically gauge the relevance of a study by briefly scanning the title and abstract. If these lack essential keywords or are mired in uncommon jargon, they may not capture the reader’s interest. An abstract that is well-structured, clear, accurate, and written with a narrative can significantly influence whether a study is read thoroughly, sidelined onto a reading backlog, or ignored [2,3,6–8]. Therefore, the interplay between strategic keyword inclusion and compelling abstract composition serves as a bridge between discoverability and engagement, laying the groundwork for academic impact (i.e., whether the study is read, cited, and/or used in future works).

Here, we propose recommendations to maximise the discoverability and impact of scientific articles. First, we offer a practical guide to crafting effective titles, abstracts, and keywords for articles to augment their findability in search engines (Fig. 1,2). Second, we surveyed 237 journals in the fields of ecology and evolutionary biology to evaluate how existing author guidelines may inadvertently hinder article discoverability. Third, we suggest a set of recommendations for journal editors that aim to optimise the likelihood of published works being discovered and cited. Ultimately, these recommendations aim to enhance article engagement and facilitate the synthesis of evidence.

**Figure 1:**
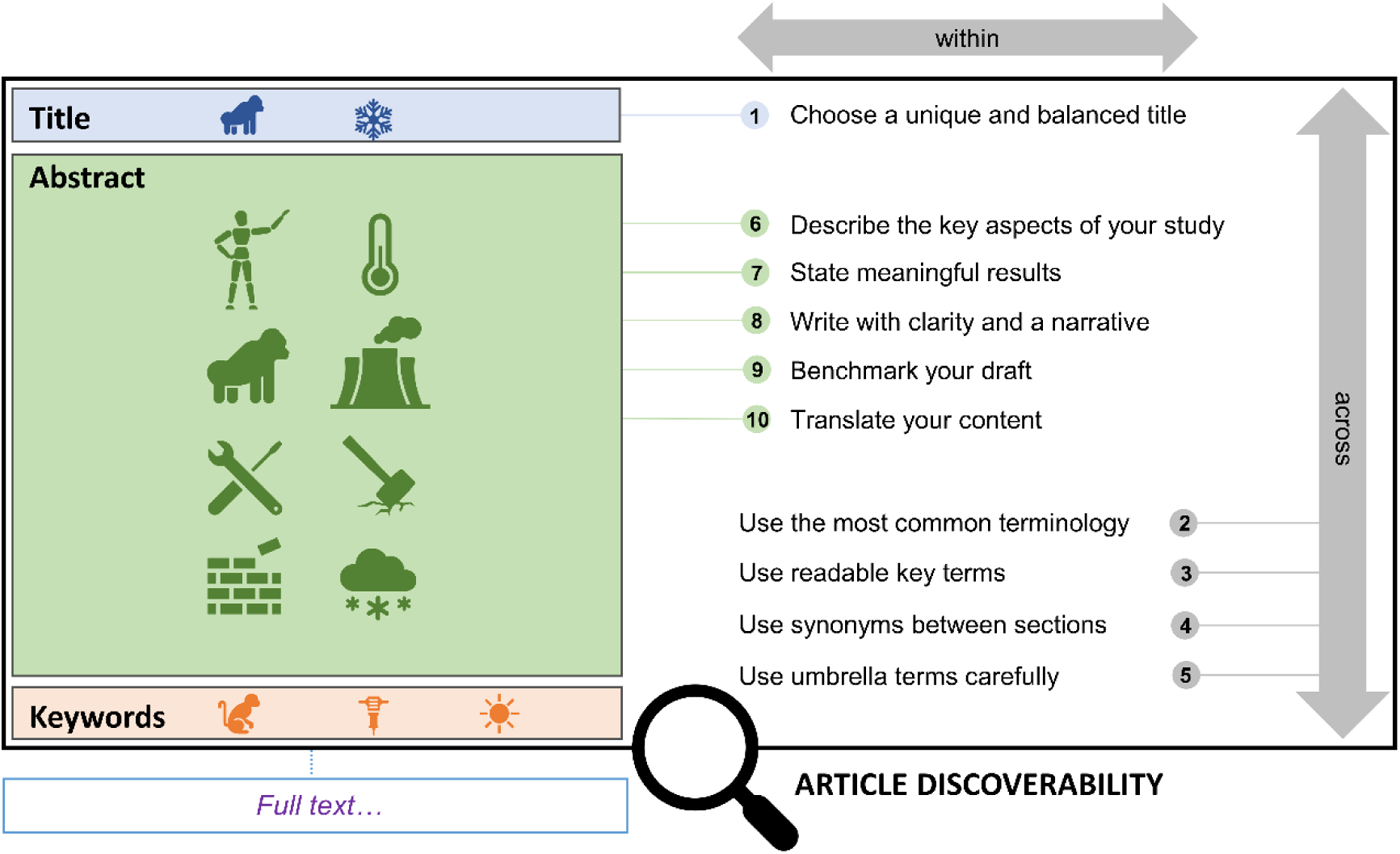
Ten strategic recommendations to improve the discoverability of scientific articles. Some recommendations are specific to a section, while others transcend multiple sections.

**Figure 2:**
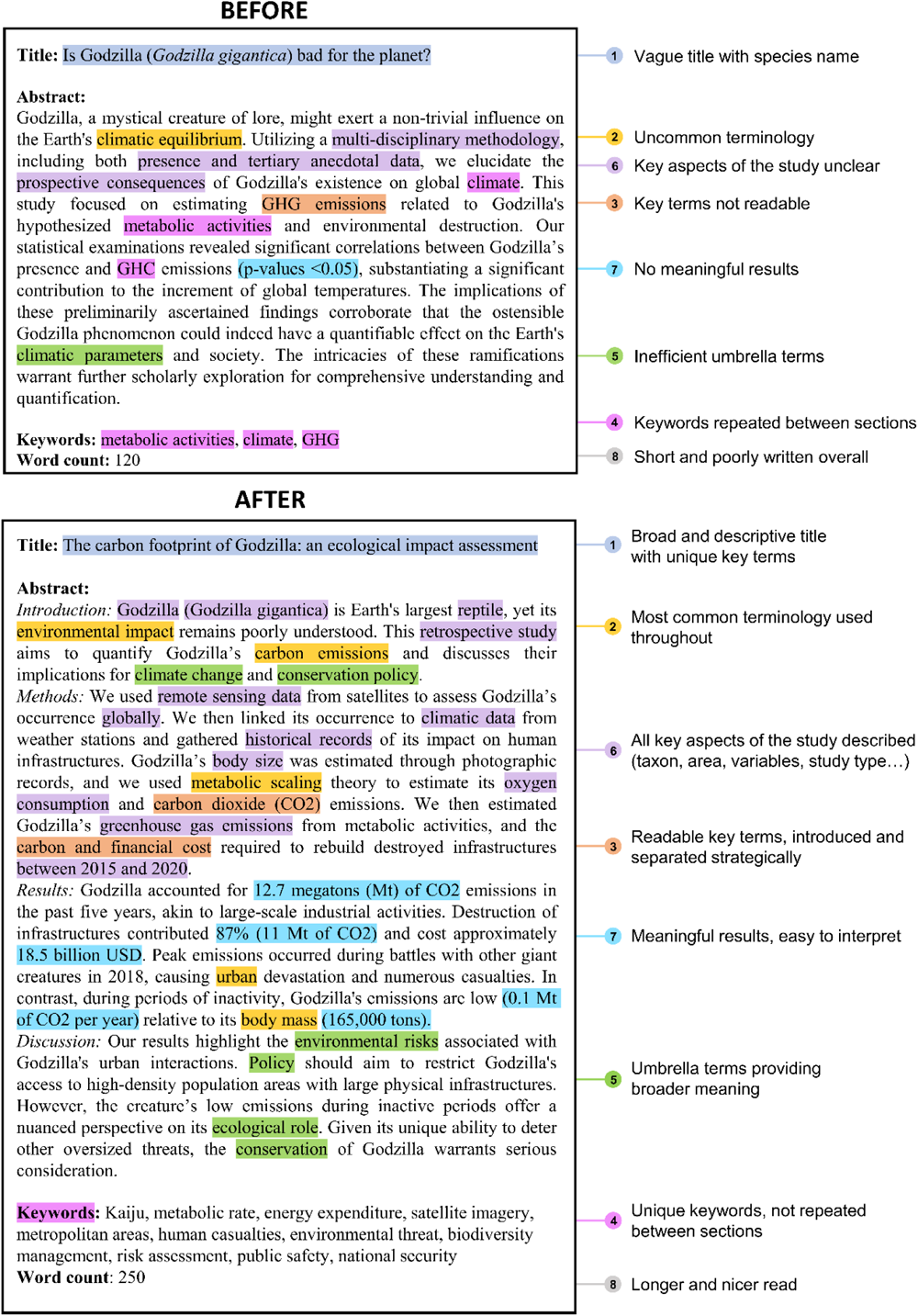
Example of strategic use of key terms. Above is a hypothetical abstract capped at 120 words that does not follow our recommendations. Below is a longer (250 words) abstract that follows our recommendations for crafting titles, abstracts, and keywords. The text highlighted and numbers refer to specific parts of the guide (see main text)

### A practical guide to crafting titles, abstracts, and keywords

Number refers to the sections of the guide (see main text).

#### 1. Choose a unique and balanced title

Titles hold a pivotal role in scientific papers. From reviewers to readers, it is the first point of engagement. It is thus not surprising that an article’s discoverability and engagement can be shaped by the contents of its title.

In the field of ecology and evolutionary biology, titles have been getting longer without much consequence on citation rates [9,10]. However, exceptionally long titles tend to fare poorly during editorial review [9]. A narrow-scoped title also tends to have negative effects, with papers that included species names in titles receiving 32% fewer citations compared to papers that avoid this practice [9]. This suggests that framing your findings in a broader context can increase your study’s appeal to readers and editors. However, it is important not to inflate the scope of your study so that the title remains accurate and informative [11,12]. For instance, a study investigating the thermal tolerance of *Pogona vitticeps* could phrase its title as “*thermal tolerance of a reptile*”, as opposed to “*thermal tolerance of reptiles*”, the latter implying the study results are applicable to all reptiles.

Humour also appears to play an important role in a paper’s future impact [10]. After controlling for journal properties, papers with titles that scored the highest for humour had nearly double the citation count as papers that received the lowest scores [10]. While incorporating humour might be perceived as a risk, evidence suggests that a well-placed pun can enrich academic writing, engaging both the intellect and the funny bone.

However, the art of crafting an engaging title is complemented by a scientific consideration for accuracy and discoverability. Implementing these stylistic findings is a progressive step, but attention must also be given to the integration of relevant key terms to accurately describe the content. A simple search of your title can also ensure its distinctiveness from other published articles, reducing the likelihood of your paper being overshadowed in the vast scientific literature.

#### 2. Use the most common terminology

The terminology used in a scientific article is not merely descriptive. Key terms can be used strategically to enhance the discoverability of scientific research and their influence extends beyond the keyword section. The more we incorporate key terms or phrases that encapsulate the essence of our research, the more likely our work is to surface in broad database searches. Emphasising recognisable key terms, those frequently employed in related literature, can significantly augment the findability of an article.

A systematic approach to choosing key terms or phrases involves scrutinising similar studies to identify the terminology predominantly used. Lexical resources or linguistic tools (e.g., Thesaurus) that provide variations of essential terms can be beneficial in this process, ensuring that a variety of relevant search terms direct readers to your work.

Avoiding ambiguity also plays a crucial role in enhancing discoverability. Precise and familiar terms often outperform their broader or less recognisable counterparts. For example, “*survival*” conveys a clearer meaning than the more expansive term “*survivorship*”, and “*bird*” resonates more readily with a broader audience than the specialised term “*avian*”.

Papers with these key terms will appear more frequently in literature searches, and hence have the potential to be more impactful.

#### 3. Use readable key terms

When crafting your abstract, it is crucial to prioritise the readers’ ability to discover and understand your work. Select key terms or phrases that are likely to appear in search queries, ensuring they are not separated by stop words or special characters that might hinder discovery. For instance, instead of “*offspring number and survival*”, consider using “*offspring number and offspring survival*” to align with typical search queries (e.g., “*offspring surviva*l”). Similarly, avoid key terms separated by hyphens or containing special characters, unless they represent the most common terminology.

Technical jargon is often difficult to circumvent in the methods section, but strive to minimise its use in the abstract. When technical terms or acronyms are necessary, choose them thoughtfully to avoid confusion. For example, “*PCR*” could stand for “*Polymerase Chain Reaction*” or “*Principal Coordinates Regression*”, potentially confusing a broad readership. In some instances, acronyms are a useful way to reduce word count and make the abstract readable, particularly for key terms that are mostly defined by their acronyms (e.g., Per- and Polyfluoroakyl Substances, PFAS). In such cases, ensure that both the acronym and its definition are in the abstract and keywords, preserving clarity while maximising discoverability.

#### 4. Use synonyms between sections

Leveraging synonyms is another tactic to maximise the chances of your article being discovered. Readers often search for relevant studies using one or few relevant key terms.

Therefore, including as many key terms across the title, abstract, and keywords will maximise the chances of a study being found [13]. To do this efficiently, consider all possible synonyms of key terms, using lexical resources, or seeking advice from field experts and collaborators. Using text-mining from samples of relevant studies is also an efficient way to find additional key terms [14].

Once your key terms are selected, strategically distribute them across your title, abstract, and keywords. For readability and clarity, maintain consistent terminology in the abstract itself, but vary verbs and adjectives to keep the writing engaging. This approach preserves the abstract’s clarity, while leveraging different synonyms in other sections to enhance discoverability. Evidence shows that articles that distributed keywords between sections had citation advantages [15]. This underscores the importance of careful keyword selection and placement, turning what might be overlooked as a minor detail into a meaningful opportunity to extend the impact of your work.

#### 5. Use umbrella terms carefully

As discussed previously, selecting the appropriate terminology in the title, abstract, and keywords is essential to accurately represent your study and reach potential readers. This involves carefully considering the use of key terms or phrases that are directly related to your research, as well as umbrella terms that can convey broader context. Umbrella terms are broad and general phrases that encompass a wide range of concepts. While they can be useful for situating your study in a larger framework and enhance discoverability, misuse or over-reliance on these terms can render your paper vague and lead to confusion [12].

For instance, if your research specifically examines the impact of deforestation on amphibian biodiversity in a particular region, it might be appropriate to mention “*biodiversity loss*” or “*environmental degradation*.” However, using overly broad terms such as “*climate change*” without direct relevance could dilute the specificity of your research, distancing it from its core audience. Similarly, using broad terms, such as “*anthropogenic impacts*”, is not optimal if the study is focusing on urban ecology or eutrophication. In navigating umbrella terms, you must strike a delicate balance between providing broader context and maintaining the specificity of your study. This will ensure that your study will be discovered by a broad audience that includes specialists and researchers from other fields.

#### 6. Describe the key aspects of your study

A recommended approach adopted by some ecological and evolutionary biology journals is to structure the abstract using the IMRAD framework (Introduction, Methods, Results, and Discussion) or derivatives. While not all journals allow structured abstracts, any abstract can be organised logically. The IMRAD framework facilitates a logical flow of information and ensures that the abstract is a stand-alone summary of the paper. It also ensures that researchers can efficiently locate specific abstract sections and gather the necessary information.

Within these abstract sections, key elements should be incorporated to enhance discoverability in online bibliographic databases. These include the taxonomic group, species name, response variable(s), independent variable(s), study area, and study type. By including these components, the abstract becomes more discoverable to researchers searching for a specific aspect of your study. For example, one may be interested in compiling studies on bird wing length in tropical birds and may search for the key terms “*birds*”, “*wing length*” and “*tropical*”. Relevant studies using alternative key terms such as “*Passerines*” instead of “*birds*”, general terms such as “*body size*” instead of “*wing length*”, or failing to include key terms such as “*tropical*”, may not be found. Optimally, all these keywords should be present to maximise your chances of being discovered.

However, deciding on the level of classification when describing these key sections is not trivial. For instance, taxonomic groups could be divided in formal levels of classification (order, family, genus, etc.), or more colloquially (birds, reptiles, fish). Study areas can also vary in granularity, ranging from the continent level to the local county. We recommend proofreading your abstract with a focus on discoverability. Imagine yourself looking for a similar study, and consider what elements of the abstract would facilitate its discovery. You may also search for similar studies, or review systematic reviews and meta-analyses on related topics. Choosing the right terminology is a compromise between discoverability and specificity, and the choice of these terms should align with the scope and audience for your study.

#### 7. State meaningful results

The presentation of results in the abstract is essential as it highlights the study’s central quantitative findings. To effectively communicate these findings, the results should be summarised concisely in one to three clear sentences, emphasising key points, and avoiding complex statistical details that might require specialised knowledge or extensive contextualisation. The results should be accessible to a wide audience, especially in ecology and evolutionary biology, where readers have varying backgrounds and expertise levels.

While null hypothesis significance testing results are commonly reported in abstracts, they do not provide information on the magnitude or practical importance of the observed effect [16,17]. Instead, the focus should be on the effect size, a measure that conveys the magnitude, direction, and precision of an effect. Effect sizes describe more meaningful information about the biological effect than statistical significance alone [18,19].

Consider, for example, a study on the impact of temperature changes on fish body size. Instead of reporting p-values or model coefficients, it would be more effective to say, “We found that a 3°C increase in water temperature led to a 15% (± 2% SD) decrease in body length”. This statement conveys the core finding with a focus on the magnitude and direction of the effect, making it more comprehensible to a wide readership without extensive contextualisation.

#### 8. Write with clarity and a narrative

The quality of an abstract is an important factor in determining the life and legacy of a paper, and requires a careful balance of accuracy, clarity, and style. Though often overlooked, adding a narrative to your abstract can elevate its appeal. While the content must be scientifically rigorous, a well-phrased abstract can make the reading experience more engaging without sacrificing scholarly value. In fact, narratives are inherently persuasive and favour engagement with a broader audience [7]. By weaving a clear narrative and connecting ideas, you can enhance both the readability and appeal of your work [6–8]. While narrativity is subjective, narrative indicators (i.e., metrics measuring the degree of narrativity) can be used to assess and develop your narrative, and are positively correlated with journal impact factors and citation rates [6].

#### 9. Benchmark your draft

A robust strategy to gauge the coverage of your key terms is to compare them with the content of similar studies. To do this, you can use your key terms in database searches (e.g., Web of Science, Scopus, or Google Scholar) to inspect their effectiveness in capturing related papers on the subject you are investigating [20]. Conversely, you can use search terms of existing systematic reviews or meta-analyses relevant to your study topic to ensure that your paper would be retrieved based on your abstract and keywords. You can do a similar exercise by attempting to do a comprehensive search of your own to think of distinct terms you can include as keywords.

Remember to recognise that key terms contained in your abstract should not be duplicated as keywords. That is because the abstract content is indexed by most databases, meaning that the words in the abstract naturally act as keywords. At this stage, it is valuable to share your abstract and keywords with co-authors to seek their insights. Sharing your draft with someone outside of your field may also help find terms that are overly technical for a broad readership. This collaborative exercise is useful to ensure you use the most relevant key terms, increasing discoverability.

#### 10. Translate your content

English is considered the *lingua franca* of scientific research, allowing it to have a global reach. However, not all scientists or readers have a good understanding of English, limiting the accessibility of vital research [21]. Recognising this, some journals allow titles and abstracts in multiple languages, although only 17% of journals in biological sciences currently offer this option [22].

Translating titles and abstracts enhances inclusivity, broadens the scope and impact of research, and serves as a bridge to overcome language barriers [22–25]. The reach of scientific studies can be expanded to include scientists, practitioners, policymakers, and the general public in non-English speaking regions by making content available in different languages when permitted [21–26]. This fosters a more balanced global understanding of scientific advancements. Furthermore, translating content can lead to broader recognition, more citations, and potential collaborations, amplifying the global resonance of scientific studies. Such a practice promotes equitable engagement in the scientific community, thereby increasing the visibility and impact of research.

##### Further considerations

Finding relevant literature can be difficult, but so is crafting an informative title, abstract, and keywords. Above, we argued that envisioning yourself conducting a literature search can lead you to become a better craftsman of the searchable face of your own work. You can also learn a lot from the methods used for conducting systematic searches of literature (e.g., see [4] for a guide for ecologists and evolutionary biologists) – especially on how search keywords are selected, and search strings are composed to find evidence used in meta-analyses and quantitative evidence syntheses. In addition, professional courses or advice from librarians are a great way to gather knowledge on the workings of search engines and systematic review searches. By understanding the strengths and limitations of search engines and human minds, you will increase the chances of your scientific contributions being noticed and used to inform policies, practices, and scientific progress.

It is also important to choose the right journal for your paper to maximise its chances of discovery. Targeting the appropriate audience, rather than merely the journal’s impact factor, should be the primary consideration. Exploring various open access options, including green open access (e.g., via preprints like BioRxiv or EcoEvoRxiv), can significantly increase the visibility and accessibility of your work [27,28]. Indeed, open-access publications have citation advantages over non-open-access counterparts in ecological journals [29,30]. In addition, advertising studies on social media can boost your study’s engagement. In fact, Twitter (now X) activities predict an article’s citation performance better than a journal’s impact factor [31,32]. Although citation performance does not directly relate to discoverability, discoverability is a prerequisite for being cited among the sea of scientific literature.

Finally, we remind readers to cite writing guides (including this one!) if they find them useful. Citing writing guides in the methods or acknowledgements section not only credits the authors, but also enhances the discoverability and usage of such guidelines. As more people embrace these recommendations, the scientific community at large will benefit from more searchable, clear, and engaging literature.

### Journal policies in the ecological and evolutionary biology literature

Above, we outlined ten strategic recommendations to optimise the most prominent marketing elements of your study. Nevertheless, these strategies must navigate the limitations imposed by journal guidelines, specifically regarding constraints on the title, abstract, and keywords. To investigate these constraints, we conducted a literature survey examining the word limits of 237 journals in the fields of ecology and evolutionary biology.

#### Methods

We report our methods as per the MeRIT guidelines [33]. On 2022/09/30, CW surveyed journals classified as “*Ecology*” or “*Evolutionary Biology*” by Clarivate Journal Citation Reports, and PPottier supplemented this list with 13 multidisciplinary journals including Nature, Nature Communications, Nature Climate Change, Scientific Reports, Science, Science Advances, Communications Biology, Proceedings of the National Academy of Sciences, Plos Biology, Biological Reviews, Current Biology, eLife, and Philosophical Transactions of the Royal Society B - Biological Sciences. Our aim was to compile a representative, though non-exhaustive, list of journals publishing studies in ecology and evolutionary biology.

To gauge recommended word limits, PPottier, ML, SB, SMD, ELM, ARM, KM, LR, JT, CW, YY and SN inspected the author guidelines of each journal as of 2022/11/28, quantifying the constraints on title and abstract length, and the maximum number of keywords permitted for standard research articles. Where a range was provided, we used the upper limit. In addition, we assessed whether the abstract layout was flexible or structured.

We further quantified the actual length of titles, abstracts, and keywords from a sample of 25 studies from each journal. In the case of multidisciplinary journals, PPottier, ML, SB, SMD, ARM, KM, PPollo, LR, JT, YY, CW and SN manually inspected studies published in 2022 in Web of Science, selecting the 25 latest ecological and evolutionary biology studies as of 2023/02/06. For other journals, PPottier conducted a range of bibliographic searches (Supporting Information S1) on 2023/09/11 in Web of Science (core collection) using The University of New South Wales’ subscription and selected 25 studies per journal for analysis.

Subsequently, PPottier processed the data using R statistical software [34] (version 4.3.0). PPottier used text mining to measure the length of titles, abstracts, and the number of keywords using the *stringr* package [35] (version 1.5.0). Note that PPottier excluded abstracts under 50 words, as they were classified as comments or opinion pieces. Furthermore, PPottier analysed whether keywords were duplicated in the title or abstract. In making comparisons between author guidelines and study samples, PPottier excluded journals without explicit word limits for titles, abstracts, or keywords. PPottier also conducted a linear regression to correlate title word length with character length, and employed predictions from this model to convert character limits to word limits. In total, we obtained a sample of 5842 studies.

#### Journal guidelines

Journal guidelines on title, abstract, and keyword limits varied greatly, with a range of 120 – 500 words and an average of 265.6 words (+/- 67.3 SD; Fig. 3A). Most commonly (31.2%), journals adhered to an abstract limit of 250 words; and nearly a quarter (22.8%) of journals did not stipulate an abstract length limit in their guidelines. Additionally, 13.1% of journals permitted structured abstracts using the IMRAD framework or derivatives.

**Figure 3:**
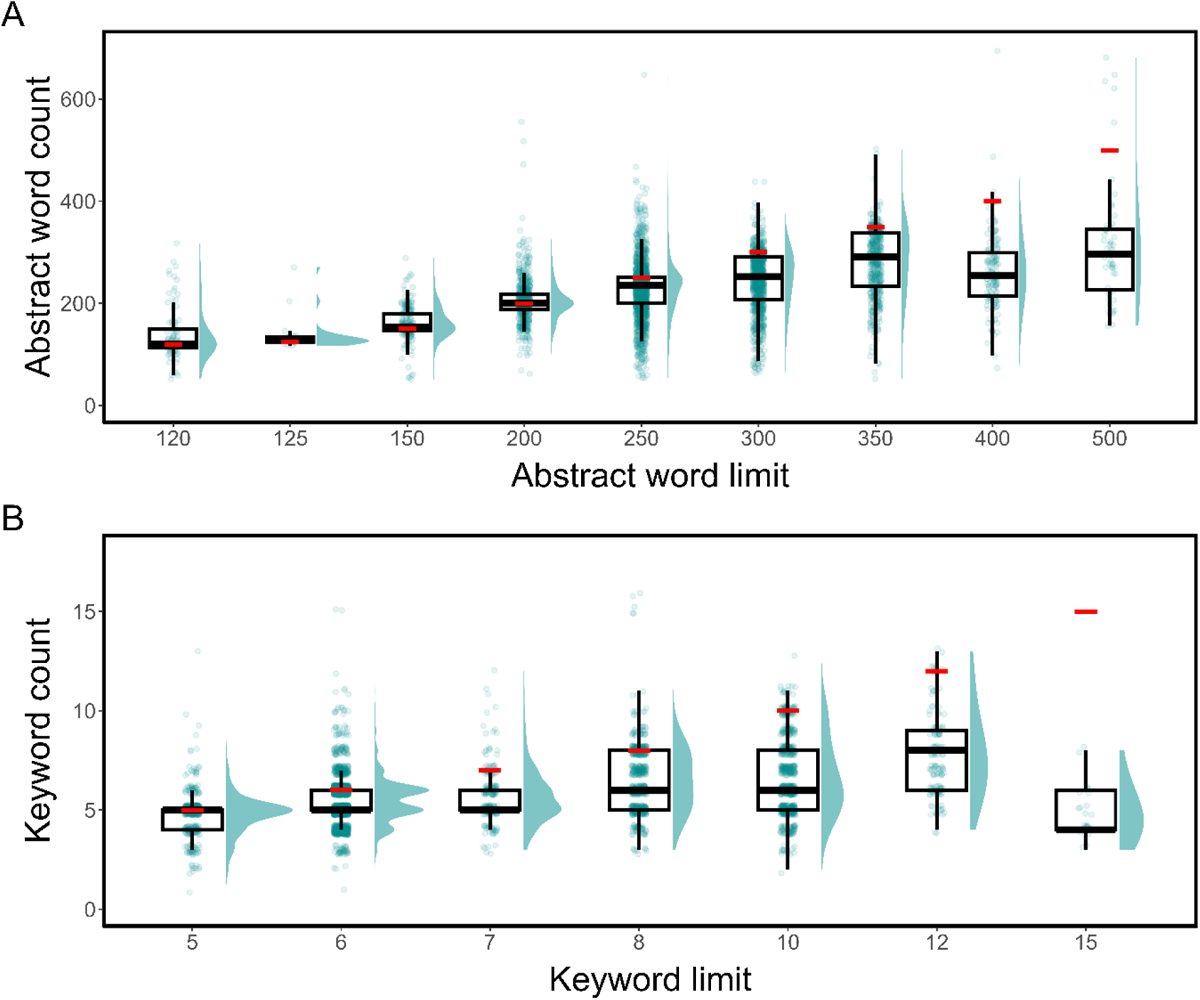
Comparison between word limits imposed by journals and the length of abstracts (A) and titles (B) in a sample of 5842 studies in ecology and evolutionary biology. Individual data points refer to abstract word or keyword counts from a sample of studies, along with their density distribution. Medians are represented by thick the black lines, interquartile ranges by the boxes, and whiskers extend to 1.5 times the interquartile range. Red lines indicate the abstract or keyword limit imposed by journals.

Titles were frequently unregulated in author guidelines, as 77.2% of journals refrained from imposing a word or character limit, at least in the author guidelines. For journals that did state word limits, titles were recommended not to exceed 15.8 words (+/- 5.2 SD) on average (range: 6 – 33).

With regards to keywords, nearly a quarter (24.1%) of journals did not mention a limit, although it must be noted that these, as well as abstracts and keywords, may be constrained within the journal’s submission platform. In cases where journals did provide a keyword limit in their guidelines, the majority capped the number of keywords at six (35.4%), with a range of 5 – 15 words and an average of 7.3 (+/- 2.0 SD; Fig. 3B).

#### Study samples

Using a sample of 25 recent studies from each journal, we found that the range of abstract lengths varied significantly, spanning from 51 to 1390 words and averaging 234.0 words (+/- 83.4 SD; n = 4441; Fig. 3A). Interestingly, abstract lengths often reflected restrictions set by journal guidelines; particularly for journals allowing less than 300 words

(Fig. 3A). For journals with word limits equal or above 300 words, abstracts were generally shorter than the word limit (Fig. 3A). It is important to acknowledge the imperfections of text mining from the databases of academic bibliographic records, and the challenges inherent to processing data exported from the Web of Science. For instance, research digests or highlights are sometimes misconstrued as the study’s abstract, thereby inflating the actual word count. Moreover, we were not able to distinguish study types (e.g., opinion pieces, commentaries, reviews), which often differ in allowed word count from standard research articles. Nevertheless, analyses of a representative sample of studies provides a useful overview of the general pattern, with outliers typically being exceptions. Indeed, we found that the median abstract length of each journal closely aligns with the guidelines (Fig. 1).

In journals stating explicit title word limits, title lengths ranged from 3 to 37 words, with an average of 13.8 words (+/- 4.4 SD; n = 1284). In other journals, title lengths were similar, averaging 14.7 words (+/- 4.5 SD; n = 4558).

In our sample of studies, we found that the number of keywords averaged 5.9 (+/- 1.6 SD; range 1 – 18; n = 4235; Fig. 3B). This figure is surprisingly lower than what author guidelines typically prescribe (7.3 keywords +/- 2.0 SD, Fig. 3B). Interestingly, 91.9% of the studies in our survey (n = 5002) duplicated at least one keyword in either the title or abstract, which we found was occasionally recommended in journal policies. On average, 2.65 (+/-1.60 SD) keywords were reused in the title or abstract, which represents 45.5% (+/- 0.25 SD) of the keywords.

### Recommendations to editors

In an era where information is increasingly digitised, publishing constraints traditionally imposed by print media may no longer be fitting. Our investigation into journal guidelines, and how authors engage with abstract length and keyword limitations, yields insights that call for a potential re-evaluation of current practices.

#### 1. Adopting structured abstracts and reconsidering word limits

Our survey reveals that authors frequently push their abstracts to the maximum allowable length (Fig. 3A). This trend is especially pronounced in journals with stringent word constraints, indicating that current word limits may be overly restrictive. Historically, these limits were rooted in the physical space constrained by printed journals. In today’s digital landscape, such limitations are less relevant and may hinder the discoverability and citation of research.

We encourage editors to consider adopting (optional) structured abstracts, which often have the advantage of ensuring that authors do not omit specifying key aspects of their study [36,37]. In the field of ecology and evolutionary biology, information such as the taxonomic group, species name, location, study type, and variables investigated are essential study aspects that should always be stated. Given that structured abstracts are typically longer [38] and that authors already approach word limits, editors may need to consider relaxing word count constraints. As demonstrated with examples using abstract lengths of 120 and 250 words (Fig. 2), an increase in word count can allow authors to supplement their abstract with additional key terms. This adjustment could significantly enhance the discoverability of studies, not only for regular author searches but also for systematic reviews.

#### 2. Optimising keyword usage

Most journals limit the number of keywords, constraining the study’s association with synonyms and relevant terms. We recommend that journals implement a large keyword limit to enhance discoverability. Increasing keyword limits is also perhaps easier to implement than increased abstract lengths. In fact, we believe there are no clear incentives to restrict the number of keywords, and both authors and journals could benefit from increased discoverability. Implementing a standardised term system that is machine readable, akin to MeSH terms for biology, could also help authors choosing the right terminology and increase indexing. Our survey also revealed that authors generally do not use all the keywords allowed. We argue that this is a missed opportunity. We encourage authors, editors, and reviewers to leverage the potential of strategic and comprehensive keyword selection.

A concerted effort by editors and reviewers to assess keywords (as well as key terms in the abstract and title) for relevance, accuracy, and redundancy can further ensure that these terms genuinely reflect the study’s content and optimise discoverability. In fact, nearly 92% of the studies we surveyed used redundant key terms between the title, abstract, and keyword sections. Ensuring that the right keywords are used and placed thoughtfully in these critical places of a study would be an important step to increase the discoverability and impact of scientific publications.

#### 3. Accepting multilingual content

Publishing multilingual abstracts and titles could significantly amplify the global resonance of scientific studies, enhancing accessibility across linguistic barriers. However, only 17% of journals in biological sciences allow multilingual abstracts [22]. We encourage editors to consider publishing multilingual summaries to increase the accessibility of scientific knowledge in countries where English is not the primary language, potentially yielding a greater impact [22–26]. For instance, FEMS (Federation of European Microbiological Societies) journals have translated the abstracts and titles of numerous articles in Portuguese and Spanish, which has significantly increased knowledge discovery (https://academic.oup.com/fems-journals/pages/alam_2018; accessed on 2023/08/23). As translation tools have yet to properly incorporate the highly specific and complex terminology in scientific research, we believe it is valuable to allow authors to submit multilingual content. By embracing multilingual abstracts and titles, editors can foster greater inclusivity and bridge the language divide, enriching the global scientific dialogue and allowing valuable research to reach an even wider audience.

## Conclusions

Crafting a title, abstract, and choosing the right keywords is an art in itself. By understanding how scientific studies are indexed in databases and searched by authors, we can strategically increase the discoverability and impact of scientific research. Particularly, the strategic use and placement of keywords can maximise indexing, in turn laying the groundwork for discoverability and impact. Comparing author guidelines with samples of studies from journals in ecology and evolutionary biology, we found that authors often push their abstract to the maximum word limit allowed, and that the number of keywords used is low and mostly redundant. These reflect restrictive guidelines that may be relics of the print era when physical page limits existed, and are not optimised to increasing the discoverability of studies in the rapidly expanding landscape of digital publications. Therefore, we encourage journals to use effective strategies to maximise the impact of their publications. By embracing these recommendations, editors can create an environment that aligns with the digital era and promotes the broader dissemination and impact of scientific research. Such actions reflect a recognition of the evolving needs of the scientific community and the critical role of discoverability in shaping the scientific knowledge landscape.

## Acknowledgements

This work has been conducted during weekly meetings with the Interdisciplinary Ecology and Evolution Lab (i-deel) at the University of New South Wales. We acknowledge the Bedegal people, the traditional custodians of the land in which this work took place. This work was not supported by a specific grant.

## Authors’ contributions

Conceptualization: PPottier, ML, SB, SMD, ELM, ARM, KM, LR, JT, CW, YY, SN. Methodology: PPottier, ML, SB, SMD, PAD, ELM, ARM, AM, KM, PPollo, LR, JT, CW,

YY, SN

Software: PPottier

Validation: PPottier

Formal analysis: PPottier

Investigation: PPottier, ML, SB, SMD, ELM, ARM, KM, PPollo, LR, JT, CW, YY, SN Resources: None.

Data curation: PPottier

Writing - Original Draft: PPottier, ML, SB, ARM, AM, KM, PPollo, LR, JT, CW, YY, SN Writing - Review & Editing: All authors

Visualization: PPottier, ML, CW

Supervision: SN

Project administration: PPottier

Funding acquisition: None.

## Data availability statement

Data, code, and additional materials are available to review at https://github.com/p-pottier/keywords_to_success and will be archived in Zenodo upon acceptance.

## Declaration of AI use

The authors declare having used GPT 4.0 and GPT 3.5 (OpenAI) to improve the clarity and readability of this work. After using these tools, the authors reviewed and edited the content as needed and take full responsibility for the content of the publication.

## References

1. Bornmann L, Mutz R. 2015 Growth rates of modern science: A bibliometric analysis based on the number of publications and cited references. Journal of the Association for Information Science and Technology 66, 2215–2222. (doi:10.1002/asi.23329)

2. Mack C. 2012 How to write a good scientific paper: title, abstract, and keywords. Journal of Micro/Nanolithography, MEMS, and MOEMS 11, 020101–020101.

3. Cook DA, Bordage G. 2016 Twelve tips on writing abstracts and titles: How to get people to use and cite your work. Medical Teacher 38, 1100–1104. (doi:10.1080/0142159X.2016.1181732)

4. Foo YZ, O’Dea RE, Koricheva J, Nakagawa S, Lagisz M. 2021 A practical guide to question formation, systematic searching and study screening for literature reviews in ecology and evolution. Methods in Ecology and Evolution 12, 1705–1720. (doi:10.1111/2041-210X.13654)

5. Hennessy EA et al. 2022 Ensuring Prevention Science Research is Synthesis-Ready for Immediate and Lasting Scientific Impact. Prevention Science 23, 809–820. (doi:10.1007/s11121-021-01279-8)

6. Hillier A, Kelly RP, Klinger T. 2016 Narrative Style Influences Citation Frequency in Climate Change Science. PLOS One 11, e0167983. (doi:10.1371/journal.pone.0167983)

7. Dahlstrom MF. 2014 Using narratives and storytelling to communicate science with nonexpert audiences. Proceedings of the National Academy of Sciences 111, 13614– 13620. (doi:10.1073/pnas.1320645111)

8. Kelly RP, Cooley SR, Klinger T. 2014 Narratives Can Motivate Environmental Action: The Whiskey Creek Ocean Acidification Story. Ambio 43, 592–599. (doi:10.1007/s13280-013-0442-2)

9. Fox CW, Burns CS. 2015 The relationship between manuscript title structure and success: editorial decisions and citation performance for an ecological journal. Ecology and Evolution 5, 1970–1980. (doi:10.1002/ece3.1480)

10. Heard SB, Cull CA, White ER. 2023 If this title is funny, will you cite me? Citation impacts of humour and other features of article titles in ecology and evolution. FACETS 8, 1–15. (doi:10.1139/facets-2022-0079)

11. Doubleday ZA, Connell SD. 2017 Publishing with Objective Charisma: Breaking Science’s Paradox. Trends in Ecology & Evolution 32, 803–805. (doi:10.1016/j.tree.2017.06.011)

12. Nature Human Behaviour. 2023 Writing more informative titles and abstracts. Nature Human Behaviour 7, 465–465. (doi:10.1038/s41562-023-01596-8)

13. Masuchika GN. 2014 Problems of scholar-created, synonymous subject terms in Buddhism. Library Review 63, 252–260. (doi:10.1108/LR-10-2013-0128)

14. Grames EM, Stillman AN, Tingley MW, Elphick CS. 2019 An automated approach to identifying search terms for systematic reviews using keyword co-occurrence networks. Methods in Ecology and Evolution 10, 1645–1654. (doi:10.1111/2041-210X.13268)

15. Rostami F, Mohammadpoorasl A, Hajizadeh M. 2014 The effect of characteristics of title on citation rates of articles. Scientometrics 98, 2007–2010. (doi:10.1007/s11192-013-1118-1)

16. Nakagawa S, Cuthill IC. 2007 Effect size, confidence interval and statistical significance: a practical guide for biologists. Biological Reviews 82, 591–605. (doi:10.1111/j.1469-185X.2007.00027.x)

17. Wasserstein RL, Schirm AL, Lazar NA. 2019 Moving to a World Beyond “p < 0.05”. The American Statistician 73, 1–19. (doi:10.1080/00031305.2019.1583913)

18. Cohen J. 1992 Things I have learned (so far). In Methodological issues & strategies in clinical research, pp. 315–333. Washington, DC, US: American Psychological Association. (doi:10.1037/10109-028)

19. Cohen J. 1994 The earth is round (p < .05). American Psychologist 49, 997–1003. (doi:10.1037/0003-066X.49.12.997)

20. Ruffell D. 2019 Writing a great abstract: tips from an Editor. FEBS Letters 593, 141–143. (doi:10.1002/1873-3468.13304)

21. Amano T et al. 2023 The manifold costs of being a non-native English speaker in science. PLOS Biology 21, e3002184. (doi:10.1371/journal.pbio.3002184)

22. Arenas-Castro H et al. 2023 Academic publishing requires linguistically inclusive policies.

23. Amano T et al. 2021 Tapping into non-English-language science for the conservation of global biodiversity. PLOS Biology 19, e3001296. (doi:10.1371/journal.pbio.3001296)

24. Nolde-Lopez B, Bundus J, Arenas-Castro H, Román D, Chowdhury S, Amano T, Berdejo-Espinola V, Wadgymar SM. 2023 Language Barriers in Organismal Biology: What Can Journals Do Better? Integrative Organismal Biology 5, obad003. (doi:10.1093/iob/obad003)

25. Amano T, González-Varo JP, Sutherland WJ. 2016 Languages Are Still a Major Barrier to Global Science. PLOS Biology 14, e2000933. (doi:10.1371/journal.pbio.2000933)

26. Amano T et al. 2023 The role of non-English-language science in informing national biodiversity assessments. Nature Sustainability 6, 845–854. (doi:10.1038/s41893-023-01087-8)

27. Harnad S, Brody T, ValliÃres F, Carr L, Hitchcock S, Gingras Y, Oppenheim C, Stamerjohanns H, Hilf ER. 2004 The Access/Impact Problem and the Green and Gold Roads to Open Access. Serials Review 30, 310–314. (doi:10.1080/00987913.2004.10764930)

28. Laakso M. 2014 Green open access policies of scholarly journal publishers: a study of what, when, and where self-archiving is allowed. Scientometrics 99, 475–494. (doi:10.1007/s11192-013-1205-3)

29. Tang M, Bever JD, Yu F-H. 2017 Open access increases citations of papers in ecology. Ecosphere 8, e01887. (doi:10.1002/ecs2.1887)

30. Clements JC. 2017 Open access articles receive more citations in hybrid marine ecology journals. FACETS 2, 1–14. (doi:10.1139/facets-2016-0032)

31. Lamb CT, Gilbert SL, Ford AT. 2018 Tweet success? Scientific communication correlates with increased citations in Ecology and Conservation. PeerJ 6, e4564. (doi:10.7717/peerj.4564)

32. Peoples BK, Midway SR, Sackett D, Lynch A, Cooney PB. 2016 Twitter Predicts Citation Rates of Ecological Research. PLOS One 11, e0166570. (doi:10.1371/journal.pone.0166570)

33. Nakagawa S et al. 2023 Method Reporting with Initials for Transparency (MeRIT) promotes more granularity and accountability for author contributions. Nature Communications 14, 1788. (doi:10.1038/s41467-023-37039-1)

34. R Core Team. 2013 R: A language and environment for statistical computing.

35. Wickham H. 2010 stringr: modern, consistent string processing. R J. 2, 38.

36. Haynes RB, Mulrow CD, Huth EJ, Altman DG, Gardner MJ. 1990 More Informative Abstracts Revisited. Annals of Internal Medicine 113, 69–76. (doi:10.7326/0003-4819-113-1-69)

37. Sharma S, Harrison JE. 2006 Structured abstracts: Do they improve the quality of information in abstracts? American Journal of Orthodontics and Dentofacial Orthopedics 130, 523–530. (doi:10.1016/j.ajodo.2005.10.023)

38. Hartley J. 2014 Current findings from research on structured abstracts: an update. Journal of the Medical Library Association 102, 146–148. (doi:10.3163/1536-5050.102.3.002)

