## Supporting Information S1 for "Keywords to success: a practical guide to maximise the visibility and impact of academic papers"

The search strings used to perform the bibliographic searches are described below. In total, seven search strings were used. All searches were performed in Web of Science (core collection) using The University of New South Wales' university subscription, on 2023/09/11. Details on how these search strings were constructed can be found at https://github.com/p-pottier/keywords_to_success

***First string (October to December 2022): 5785 results***

IS=("0378-1844" OR "0717-6317" OR "0973-7308" OR "1029-0370" OR "1076-836X" OR "1095-8312" OR "1095-8606" OR "1095-922X" OR "1095-9513" OR "1096-0031" OR "1096-0325" OR "1096-8644" OR "1176-9343" OR "1177-7788" OR "1314-2488" OR "1338-7014" OR "1348-8570" OR "1365-2028" OR "1365-2427" OR "1365-2435" OR "1365-2486" OR "1365-2540" OR "1365-2656" OR "1365-2664" OR "1365-2699" OR "1365-2745" OR "1365-2907" OR "1365-294X" OR "1365-3008" OR "1365-3113" OR "1399-1183" OR "1420-9101" OR "1423-0445" OR "1424-2818" OR "1432-041X" OR "1432-0762" OR "1432-1432" OR "1432-184X" OR "1432-1939" OR "1432-2056" OR "1433-8319" OR "1435-0629" OR "1438-390X" OR "1439-0469" OR "1439-0574" OR "1440-1703" OR "1442-1984" OR "1442-8903" OR "1442-9993" OR "1446-5701" OR "1447-2600" OR "1448-5494" OR "1461-0248" OR "1463-6409" OR "1464-5262" OR "1465-7279" OR "1465-7333" OR "1466-8238" OR "1469-1795" OR "1469-7831" OR "1471-2148" OR "1471-2954" OR "1472-4642" OR "1472-6785" OR "1475-3057" OR "1476-9840" OR "1478-0941" OR "1523-1739" OR "1525-142X" OR "1526-100X" OR "1537-1719" OR "1537-5323" OR "1540-9309" OR "1543-4079" OR "1545-2069" OR "1548-2324" OR "1551-5028" OR "1552-5015" OR "1557-7015" OR "1558-5646" OR "1572-9710" OR "1572-9761" OR "1573-1464" OR "1573-1561" OR "1573-1642" OR "1573-3017" OR "1573-5052" OR "1573-5125" OR "1573-5133" OR "1573-8477" OR "1573-8477" OR "1588-2756" OR "1600-0587" OR "1600-0706" OR "1608-3202" OR "1608-3334" OR "1615-6110" OR "1616-1564" OR "1616-1599" OR "1618-0089" OR "1618-0585" OR "1618-1077" OR "1618-1093" OR "1654-109X" OR "1654-1103" OR "1726-4189" OR "1727-9380" OR "1744-7429" OR "1744-957X" OR "1745-1019" OR "1745-2627" OR "1751-3766" OR "1751-7370" OR "1751-8369" OR "1752-4571" OR "1752-993X" OR "1755-0998" OR "1758-6798" OR "1759-6653" OR "1778-3615" OR "1785-0037" OR "1797-2450" OR "1809-4392" OR "1818-5487" OR "1834-7541" OR "1860-188X" OR "1872-6062" OR "1872-6992" OR "1872-7026" OR "1872-8383" OR "1873-1511" OR "1873-2305" OR "1873-2917" OR "1873-2925" OR "1873-6106" OR "1873-6238" OR "1874-1746" OR "1876-312X" OR "1878-0083" OR "1878-0512" OR "1879-0445" OR "1879-1697" OR "1882-5729" OR "1903-220X" OR "1918-3178" OR "1933-9747" OR "1934-2845" OR "1936-0592" OR "1936-8046" OR "1937-2817" OR "1937-3546" OR "1938-4238" OR "1938-5307" OR "1938-5331" OR "1938-5331" OR "1938-5412" OR "1938-5455" OR "1939-5582" OR "1939-9170" OR "1941-3300" OR "1943-6246" OR "1943-6262" OR "1944-8341" OR "1993-9507" OR "1995-4263" OR "1996-8175" OR "2041-2096" OR "2041-2851" OR "2041-9139" OR "2045-7758" OR "2045-7758" OR "2050-3385" OR "2050-6201" OR "2051-1434"

OR "2051-3933" OR "2056-3485" OR "2056-3744" OR "2073-1558" OR "2075-1125" OR "2079-0988" OR "2080-3397" OR "2081-8262" OR "2083-5469" OR "2150-8925" OR "2156-6941" OR "2161-9565" OR "2161-9859" OR "2162-4135" OR "2162-4399" OR "2163-582X" OR "2192-1709" OR "2212-0416" OR "2213-2244" OR "2214-5753" OR "2224-4662" OR "2296-701X" OR "2311-2077" OR "2326-2397" OR "2332-8878" OR "2348-8980" OR "2351-9894" OR "2352-2496" OR "2352-4855" OR "2363-7153" OR "2368-7460" OR "2376-6808" OR "2376-7626" OR "2397-334X" OR "2410-8200" OR "2413-0958" OR "2517-4843" OR "2520-2529" OR "2571-6255" OR "2572-2611" OR "2575-8314" OR "2588-3526" OR "2618-8406" OR "2624-893X" OR "2661-8982" OR "2662-2297" OR "2730-7182" OR "1708-3087" OR "0097-3157" OR "1944-687X" OR "1505-2249" OR "0240-8759" OR "0370-047X" OR "0028-0712" OR "1697-2473") AND LD=(2022-10-01/2022-12-25)

***Second string (June – December 2022): 4120 results***

IS=("1809-4392" OR "1873-6238" OR "1365-2028" OR "1727-9380" OR "2348-8980" OR "1938-4238" OR "2050-3385" OR "1469-1795" OR "1797-2450" OR "2041-2851" OR "1654-109X" OR "1573-5125" OR "1818-5487" OR "1616-1564" OR "2368-7460" OR "2079-0988" OR "1446-5701" OR "1618-0089" OR "1465-7279" OR "1095-8312" OR "1744-7429" OR "2162-4135" OR "1029-0370" OR "1423-0445" OR "1096-0031" OR "1588-2756" OR "2051-1434" OR "1995-4263" OR "1432-041X" OR "1600-0587" OR "2080-3397" OR "1476-9840" OR "1557-7015" OR "2192-1709" OR "2083-5469" OR "1440-1703" OR "1543-4079" OR "1708-3087" OR "2376-7626" OR "2332-8878" OR "1435-0629" OR "1573-3017" OR "1778-3615" OR "1439-0574" OR "2041-9139" OR "1525-142X" OR "2056-3744" OR "1176-9343" OR "1934-2845" OR "1573-8477" OR "1933-9747" OR "2352-2496" OR "2161-9565" OR "1540-9309" OR "1759-6653" OR "1365-2540" OR "1936-8046" OR "2517-4843" OR "0378-1844" OR "2213-2244" OR "1745-2627" OR "1447-2600" OR "2224-4662" OR "1751-3766" OR "1573-1561" OR "1879-1697" OR "1552-5015" OR "2156-6941" OR "1465-7333" OR "1095-8606" OR "1432-1432" OR "1464-5262" OR "1752-993X" OR "1941-3300" OR "1478-0941" OR "1469-7831" OR "1654-1103" OR "2588-3526" OR "1937-2817" OR "1439-0469" OR "1860-188X" OR "1365-2907" OR "1745-1019" OR "1755-0998" OR "2051-3933" OR "2162-4399" OR "1314-2488" OR "2376-6808" OR "1177-7788" OR "1938-5307" OR "2161-9859" OR "1618-1077" OR "1365-3008" OR "1938-5331" OR "1873-1511" OR "2575-8314" OR "1433-8319" OR "2363-7153" OR "1573-5052" OR "1442-1984" OR "1615-6110" OR "2572-2611" OR "1432-2056" OR "1475-3057" OR "1751-8369" OR "1505-2249" OR "1438-390X" OR "1551-5028" OR "1834-7541" OR "2056-3485" OR "0717-6317" OR "2075-1125" OR "2662-2297" OR "2413-0958" OR "1938-5412" OR "1943-6262" OR "1365-3113" OR "1996-8175" OR "1874-1746" OR "1096-0325" OR "1872-8383" OR "2661-8982" OR "1882-5729" OR "1573-1642" OR "1399-1183" OR "1944-8341" OR "1903-220X" OR "1448-5494" OR "1463-6409") AND LD=(2022-06-01/2022-12-25)

***Third string (Jan – December 2022): 1927 results.***

IS=("1727-9380" OR "1938-4238" OR "2050-3385" OR "1797-2450" OR "1818-5487" OR "1616-1564" OR "1446-5701" OR "2162-4135" OR "1423-0445" OR "1096-0031" OR "1588-2756" OR "1432-041X" OR "1476-9840" OR "2083-5469" OR "1543-4079" OR "2376-7626" OR "2041-9139" OR "1525-142X" OR "2056-3744" OR "1176-9343" OR "1934-2845" OR "1933-9747" OR "2352-2496" OR "2161-9565" OR "1540-9309" OR "1936-8046" OR "2224-4662" OR "1751-3766" OR "1552-5015" OR "2156-6941" OR "1432-1432" OR "1478-0941" OR "1439-0469" OR "1860-188X" OR "1365-2907" OR "2162-4399" OR "2376-6808" OR "1177-7788" OR "1938-5307" OR "2161-9859" OR "1938-5331" OR "1873-1511" OR "1433-8319" OR "2363-7153" OR "1442-1984" OR "1615-6110" OR "1751-8369" OR "1505-2249" OR "1438-390X" OR "1834-7541" OR "0717-6317" OR "2662-2297" OR "2413-0958" OR "1943-6262" OR "1365-3113" OR "1874-1746" OR "1096-0325" OR "1882-5729" OR "1399-1183" OR "1944-8341" OR "1903-220X") AND LD=(2022-01-01/2022-12-25)

***Fourth string (Jan 2021 to December 2022): 1253 results***

IS=("1797-2450" OR "1616-1564" OR "1446-5701" OR "2162-4135" OR "1423-0445" OR "1096-0031" OR "1432-041X" OR "1543-4079" OR "2376-7626" OR "2041-9139" OR "1525-142X" OR "1176-9343" OR "1936-8046" OR "2224-4662" OR "1439-0469" OR "1365-2907" OR "2161-9859" OR "2363-7153" OR "1751-8369" OR "1505-2249" OR "1834-7541" OR "0717-6317" OR "1943-6262" OR "1874-1746" OR "1882-5729" OR "1399-1183") AND LD=(2021-01-01/2022-12-25)

***Fifth string (Jan 2019 to December 2022): 247 results***

IS=("2162-4135" OR "1432-041X" OR "2363-7153" OR "0717-6317" OR "1882-5729" OR "1399-1183") AND LD=(2019-01-01/2022-12-25)

***Sixth string (Jan 2021 to December 2022): 2473 results***

IS=("1545-2069" OR “1876-312X” OR "1937-3546" OR "1348-8570" OR "1548-2324" OR "1918-3178" OR "1442-8903" OR “1338-7014” OR “2081-8262” OR “1876-4428” OR “2050-6201” OR “1076-836X” OR “2326-2397” OR “0097-3157” OR “2073-1558” OR “1608-3334” OR “0240-8759” OR “1993-9507” OR “1096-8644” OR “0973-7308” OR “2311-2077” OR “2520-2529” OR “1938-5455” OR “1944-687X” OR “0028-0712” OR “2410-8200”) AND LD=(2021-01-01/2022-12-25)

***Seventh string (Jan 2018 to December 2022): 824 results***

IS=("1545-2069" OR “1348-8570” OR "1937-3546" OR "2326-2397" OR "1918-3178" OR "0097-3157" OR “1076-836X” OR “1938-5455”) AND LD=(2018-01-01/2022-12-25)

***Search string for exporting multidisciplinary journals as bibtex: 329 results***

TI=("Animals in flow - towards the scientific study of intrinsic reward in animals") OR TI=("The ecological drivers and consequences of wildlife trade") OR TI=("The ephemeral resource patch concept") OR TI=("The kin selection theory of genomic imprinting and modes of reproduction in the eusocial Hymenoptera") OR TI=("Ecological integrity of tropical secondary forests: concepts and indicators") OR TI=("Solutions to fire and shade: resprouting, growing tall and the origin of Eurasian temperate broadleaved forest") OR TI=("Feather function and the evolution of birds") OR TI=("Testicular heat stress, a historical perspective and two postulates for why male germ cells are heat sensitive") OR TI=("The trace metal economy of the coral holobiont: supplies, demands and exchanges") OR TI=("Exercise builds the scaffold of life: muscle extracellular matrix biomarker responses to physical activity, inactivity, and aging") OR TI=("Long-term sperm storage in eusocial Hymenoptera") OR TI=("The phenotypic costs of captivity") OR TI=("Sex roles and sex ratios in animals") OR TI=("Extreme weather events threaten biodiversity and functions of river ecosystems: evidence from a meta-analysis") OR TI=("Centring individual animals to improve research and citation practices") OR TI=("The best of two worlds: ecology and evolution of ambophilous plants") OR TI=("Worms and gills, plates and spines: the evolutionary origins and incredible disparity of deuterostomes revealed by fossils, genes, and development") OR TI=("Hydrodynamics in early animal evolution") OR TI=("Animal linguistics: a primer") OR TI=("Perils of ingesting harmful prey by advanced snakes") OR TI=("The early diversification of ray-finned fishes (Actinopterygii): hypotheses, challenges and future prospects") OR TI=("Conformity in mate choice, the overlooked social component of animal and human culture") OR TI=("Evaluation of physiological stress in free-ranging bears: current knowledge and future directions") OR TI=("Tactics of evasion: strategies used by signallers to deter eavesdropping enemies from exploiting communication systems") OR TI=("A standalone incompatible insect technique enables mosquito suppression in the urban subtropics") OR TI=("Evolution and diversification of Mountain voles (Rodentia: Cricetidae)") OR TI=("Evaluation of the current understanding of the impact of climate change on coral physiology after three decades of experimental research") OR TI=("Simultaneous invasion decouples zebra mussels and water clarity") OR TI=("Significance of NatB-mediated N-terminal acetylation of auxin biosynthetic enzymes in maintaining auxin homeostasis in Arabidopsis thaliana") OR TI=("Palau's warmest reefs harbor thermally tolerant corals that thrive across different habitats") OR TI=("A new confuciusornithid bird with a secondary epiphyseal ossification reveals phylogenetic changes in confuciusornithid flight mode") OR TI=("A chromosome-level genome assembly reveals genomic characteristics of the American mink (Neogale vison)") OR TI=("Adaptations by the coral Acropora tenuis confer resilience to future thermal stress") OR TI=("Machine learning prediction of connectivity, biodiversity and resilience in the Coral Triangle") OR TI=("Establishment of a salt-induced bioremediation platform from marine Vibrio natriegens") OR TI=("Gamete dimorphism of the isogamous green alga (Chlamydomonas reinhardtii), is regulated by the mating type-determining gene, MID") OR TI=("An artificial neural network explains how bats might use vision for navigation") OR TI=("Climate, currents and species traits contribute to early stages of marine species redistribution") OR TI=("A non-avian dinosaur with a streamlined body exhibits potential adaptations for swimming") OR TI=("Temperature-robust rapid eye movement and slow wave sleep in the lizard Laudakia vulgaris") OR TI=("Current trends suggest most Asian countries are unlikely to meet future biodiversity targets on protected areas") OR TI=("The evolution of plant proton pump regulation via the R domain may have facilitated plant terrestrialization") OR TI=("Fossil bone histology reveals ancient origins for rapid juvenile growth in tetrapods") OR TI=("Parasitic infection increases risk-taking in a social, intermediate host carnivore") OR TI=("Dark wing pigmentation as a mechanism for improved flight efficiency in the Larinae") OR TI=("A calcitonin receptor-expressing subregion of the medial preoptic area is involved in alloparental tolerance in common marmosets") OR TI=("Identifying behavioral structure from deep variational embeddings of animal motion") OR TI=("A globally distributed durophagous marine reptile clade supports the rapid recovery of pelagic ecosystems after the Permo-Triassic mass extinction") OR TI=("Climate service driven adaptation may alleviate the impacts of climate change in agriculture") OR TI=("Species-specific song responses emerge as a by-product of tuning to the local dialect") OR TI=("Land-use and climate risk assessment for Earth's wilderness") OR TI=("Indigenous lands in protected areas have high forest integrity across the tropics") OR TI=("Genomic analysis reveals cryptic diversity in aphelids and sheds light on the emergence of Fungi") OR TI=("The Daphnia carapace and other novel structures evolved via the cryptic persistence of serial homologs") OR TI=("Ambitious global targets for mangrove and seagrass recovery") OR TI=("Natural and human-driven selection of a single non-coding body size variant in ancient and modern canids") OR TI=("Inflammation and convergent placenta gene co-option contributed to a novel reproductive tissue") OR TI=("The origins of the killer whale ecomorph") OR TI=("Social integration influences fitness in allied male dolphins") OR TI=("A vast icefish breeding colony discovered in the Antarctic") OR TI=("Conservation of locomotion-induced oculomotor activity through evolution in mammals") OR TI=("Report Contrasting modes of macro and microsynteny evolution in a eukaryotic subphylum") OR TI=("Report Fiddler crabs are unique in timing their escape responses based on speed-dependent visual cues") OR TI=("A three-eyed radiodont with fossilized neuroanatomy informs the origin of the arthropod head and segmentation") OR TI=("Foraging range scales with colony size in high-latitude seabirds") OR TI=("Phylogenomics of the world's otters") OR TI=("Macroevolutionary trends in theropod dinosaur feeding mechanics") OR TI=("Rapid evolution of pollen and pistil traits as a response to sexual selection in the post-pollination phase of mating") OR TI=("Running in the wild: Energetics explain ecological running speeds") OR TI=("Abdominal serial homologues of wings in Paleozoic insects") OR TI=("A Triassic tritrophic triad documents an early food-web cascade") OR TI=("Ant phylogenomics reveals a natural selection hotspot preceding the origin of complex eusociality") OR TI=("Early evolution of wing scales prior to the rise of moths and butterflies") OR TI=("Coevolution of motor cortex and behavioral specializations associated with flight and echolocation in bats") OR TI=("Behavioral signatures of structured feature detection during courtship in Drosophila") OR TI=("Wildlife trade targets colorful birds and threatens the aesthetic value of nature") OR TI=("Grouping behavior in a Triassic marine apex predator") OR TI=("Rapid growth preceded gigantism in sauropodomorph evolution") OR TI=("Climate fluctuations influence variation in group size in a cooperative bird") OR TI=("Rapid adaptation of a complex trait during experimental evolution of Mycobacterium tuberculosis") OR TI=("Admixture of evolutionary rates across a butterfly hybrid zone") OR TI=("A general decoding strategy explains the relationship between behavior and correlated variability") OR TI=("Contrasting parental roles shape sex differences in poison frog space use but not navigational performance") OR TI=("The spatiotemporal patterns of major human admixture events during the European Holocene") OR TI=("The Jurassic rise of squamates as supported by lepidosaur disparity and evolutionary rates") OR TI=("Evolutionary rescue of phosphomannomutase deficiency in yeast models of human disease") OR TI=("Global analysis of cytosine and adenine DNA modifications across the tree of life") OR TI=("Evolution of sexual conflict in scorpionflies") OR TI=("Emergence of behaviour in a self-organized living matter network") OR TI=("Circadian programming of the ellipsoid body sleep homeostat in Drosophila") OR TI=("Environmental selection overturns the decay relationship of soil prokaryotic community over geographic distance across grassland biotas") OR TI=("Late-life fitness gains and reproductive death in Cardiocondyla obscurior ants") OR TI=("Mammals adjust diel activity across gradients of urbanization") OR TI=("Early life stressful experiences escalate aggressive behavior in adulthood via changes in transthyretin expression and function") OR TI=("Heritability and cross-species comparisons of human cortical functional organization asymmetry") OR TI=("Ancestral reconstruction of duplicated signaling proteins reveals the evolution of signaling specificity") OR TI=("Humanization of wildlife gut microbiota in urban environments") OR TI=("Lung evolution in vertebrates and the water-to-land transition") OR TI=("Stress diminishes outcome but enhances response representations during instrumental learning") OR TI=("Using population selection and sequencing to characterize natural variation of starvation resistance in Caenorhabditis elegans") OR TI=("Fast bacterial growth reduces antibiotic accumulation and efficacy") OR TI=("An Archaea-specific c-type cytochrome maturation machinery is crucial for methanogenesis in Methanosarcina acetivorans") OR TI=("Directed evolution of the rRNA methylating enzyme Cfr reveals molecular basis of antibiotic resistance") OR TI=("Species-specific chromatin landscape determines how transposable elements shape genome evolution") OR TI=("The molecular evolution of spermatogenesis across mammals") OR TI=("Logged tropical forests have amplified and diverse ecosystem energetics") OR TI=("Microbial predators form a new supergroup of eukaryotes") OR TI=("Pathogen spillover driven by rapid changes in bat ecology") OR TI=("Light competition drives herbivore and nutrient effects on plant diversity") OR TI=("Small rainfall changes drive substantial changes in plant coexistence") OR TI=("Environmental signal in the evolutionary diversification of bird skeletons") OR TI=("Extreme escalation of heat failure rates in ectotherms with global warming") OR TI=("More losses than gains during one century of plant biodiversity change in Germany") OR TI=("Ion regulation at gills precedes gas exchange and the origin of vertebrates") OR TI=("Global trends of cropland phosphorus use and sustainability challenges") OR TI=("Global hotspots for soil nature conservation") OR TI=("The origin of placental mammal life histories") OR TI=("Even modest climate change may lead to major transitions in boreal forests") OR TI=("Sufficient conditions for rapid range expansion of a boreal conifer") OR TI=("Warm springs alter timing but not total growth of temperate deciduous trees") OR TI=("Direct evidence for phosphorus limitation on Amazon forest productivity") OR TI=("Competition for pollinators destabilizes plant coexistence") OR TI=("Emerging signals of declining forest resilience under climate change") OR TI=("A male steroid controls female sexual behaviour in the malaria mosquito") OR TI=("Increasing the resilience of plant immunity to a warming climate") OR TI=("Genome evolution and diversity of wild and cultivated potatoes") OR TI=("An oxygen-sensing mechanism for angiosperm adaptation to altitude") OR TI=("Enhanced silica export in a future ocean triggers global diatom decline") OR TI=("Tropical tree mortality has increased with rising atmospheric water stress") OR TI=("Floods differ in a warmer future") OR TI=("Greater evolutionary divergence of thermal limits within marine than terrestrial species") OR TI=("Younger trees in the upper canopy are more sensitive but also more resilient to drought") OR TI=("Populations adapt more to temperature in the ocean than on land") OR TI=("Climate change reshuffles northern species within their niches (vol 12, pg 587, 2022)") OR TI=("Behaviour as leverage") OR TI=("Future biological control") OR TI=("Soil and plants lose more water under drought") OR TI=("Tipping points of marine phytoplankton to multiple environmental stressors") OR TI=("Unique thermal sensitivity imposes a cold-water energetic barrier for vertical migrators") OR TI=("Global decline of pelagic fauna in a warmer ocean") OR TI=("Understanding eco-anxiety") OR TI=("Epigenetic plasticity enables copepods to cope with ocean acidification") OR TI=("Climate change impacts the vertical structure of marine ecosystem thermal ranges") OR TI=("Urban forests facing climate risks") OR TI=("Increased drought effects on the phenology of autumn leaf senescence") OR TI=("Selection counteracts developmental plasticity in body-size responses to climate change") OR TI=("A climate risk index for marine life") OR TI=("Climate change exacerbates almost two-thirds of pathogenic diseases affecting humans") OR TI=("Fall and rise of the phytoplankton") OR TI=("Vertically migrating phytoplankton fuel high oceanic primary production") OR TI=("Mangrove forests are facing challenges from global seawater density changes") OR TI=("Mangrove dispersal disrupted by projected changes in global seawater density") OR TI=("Value wild animals' carbon services to fill the biodiversity financing gap") OR TI=("Northern wildlife feels the heat") OR TI=("Parabrachial-to-parasubthalamic nucleus pathway mediates fear-induced suppression of feeding in male mice") OR TI=("Suppression of flavivirus transmission from animal hosts to mosquitoes with a mosquito-delivered vaccine") OR TI=("Biodiversity-stability relationships strengthen over time in a long-term grassland experiment") OR TI=("Evolutionary origins of the prolonged extant squamate radiation") OR TI=("Plant genetic diversity affects multiple trophic levels and trophic interactions") OR TI=("A diverse Ediacara assemblage survived under low-oxygen conditions") OR TI=("Increased fire activity under high atmospheric oxygen concentrations is compatible with the presence of forests") OR TI=("Resource sharing is sufficient for the emergence of division of labour") OR TI=("High-throughput screening of caterpillars as a platform to study host-microbe interactions and enteric immunity") OR TI=("US winter wheat yield loss attributed to compound hot-dry-windy events") OR TI=("The unknown biogeochemical impacts of drying rivers and streams") OR TI=("Wildflower phenological escape differs by continent and spring temperature") OR TI=("Genomic signatures of recent convergent transitions to social life in spiders") OR TI=("Consistent diel activity patterns of forest mammals among tropical regions") OR TI=("Biological effects of the loss of homochirality in a multicellular organism") OR TI=("Host biology, ecology and the environment influence microbial biomass and diversity in 101 marine fish species") OR TI=("Intensive grassland management disrupts below-ground multi-trophic resource transfer in response to drought") OR TI=("Ordovician opabiniid-like animals and the role of the proboscis in euarthropod head evolution") OR TI=("Two simple movement mechanisms for spatial division of labour in social insects") OR TI=("Calcium-mediated rapid movements defend against herbivorous insects in Mimosa pudica") OR TI=("A Wolbachia factor for male killing in lepidopteran insects") OR TI=("Altered developmental programs and oriented cell divisions lead to bulky bones during salamander limb regeneration") OR TI=("Influenza A virus reassortment in mammals gives rise to genetically distinct within-host subpopulations") OR TI=("Social interactions lead to motility-induced phase separation in fire ants") OR TI=("L-threonine promotes healthspan by expediting ferritin-dependent ferroptosis inhibition in C. elegans") OR TI=("Immune function of the serosa in hemimetabolous insect eggs") OR TI=("Mesothelial fusion mediates chorioallantoic membrane formation") OR TI=("Comparing the potential for maternal-fetal signalling in oviparous and viviparous lizards") OR TI=("Simple and complex, sexually dimorphic retinal mosaic of fritillary butterflies") OR TI=("EnteroBase: hierarchical clustering of 100 000s of bacterial genomes into species/subspecies and populations") OR TI=("The effect of sequencing and assembly on the inference of horizontal gene transfer on chromosomal and plasmid phylogenies") OR TI=("Genome-scale metabolic network reconstructions of diverse Escherichia strains reveal strain-specific adaptations") OR TI=("The social dynamics of complex gestural communication in great and lesser apes (Pan troglodytes, Pongo abelii, Symphalangus syndactylus)") OR TI=("Thermal imaging reveals social monitoring during social feeding in wild chimpanzees") OR TI=("Flexible use of contact calls in a species with high fission-fusion dynamics") OR TI=("Crested macaque facial movements are more intense and stereotyped in potentially risky social interactions") OR TI=("Language as a tool for social bonding: evidence from wild chimpanzee gestural, vocal and bimodal signals") OR TI=("Coevolution of social and communicative complexity in lemurs") OR TI=("Flexible signalling strategies by victims mediate post-conflict interactions in bonobos") OR TI=("Are ape gestures like words? Outstanding issues in detecting similarities and differences between human language and ape gesture") OR TI=("Intentional gesturing increases social complexity by allowing recipient's understanding of intentions when it is inhibited by stress") OR TI=("Socioecological complexity in primate groups and its cognitive correlates") OR TI=("The interaction engine: cuteness selection and the evolution of the interactional base for language") OR TI=("Great ape communication as contextual social inference: a computational modelling perspective") OR TI=("Social tolerance and interactional opportunities as drivers of gestural redoings in orangutans") OR TI=("Sequence organization and embodied mutual orientations: openings of social interactions between baboons") OR TI=("Comparing management strategies for conserving communities of climate-threatened species with a stochastic metacommunity model") OR TI=("Complexity-biodiversity relationships on marine urban structures: reintroducing habitat heterogeneity through eco-engineering") OR TI=("From coral reefs to Joshua trees: What ecological interactions teach us about the adaptive capacity of biodiversity in the Anthropocene") OR TI=("A resilience sensing system for the biosphere") OR TI=("Polygenic signals of sex differences in selection in humans from the UK Biobank") OR TI=("A sensitive and specific genetically encoded potassium ion biosensor for in vivo applications across the tree of life") OR TI=("Phage endolysins are adapted to specific hosts and are evolutionarily dynamic") OR TI=("An evolutionary conserved detoxification system for membrane lipid-derived peroxyl radicals in Gram-negative bacteria") OR TI=("Metacommunity analyses show an increase in ecological specialisation throughout the Ediacaran period") OR TI=("Further evidence for the capacity of mirror self-recognition in cleaner fish and the significance of ecologically relevant marks") OR TI=("Meta-analysis reveals an extreme "decline effect" in the impacts of ocean acidification on fish behavior") OR TI=("One fish, two fish, red fish, dead fish: Detecting the genomic footprint of ecological incompatibilities") OR TI=("Evolutionary pathways to SARS-CoV-2 resistance are opened and closed by epistasis acting on ACE2") OR TI=("Diversity, taxonomy, and evolution of archaeal viruses of the class Caudoviricetes") OR TI=("A cloud-based toolbox for the versatile environmental annotation of biodiversity data") OR TI=("Spatial subsidies drive sweet spots of tropical marine biomass production") OR TI=("Fitness costs of female choosiness are low in a socially monogamous songbird") OR TI=("A phylogenomic framework for charting the diversity and evolution of giant viruses") OR TI=("Rapid increase in snake dietary diversity and complexity following the end-Cretaceous mass extinction") OR TI=("Tapping into non-English-language science for the conservation of global biodiversity") OR TI=("Microplastics and anthropogenic fibre concentrations in lakes reflect surrounding land use") OR TI=("Plant pathogens convergently evolved to counteract redundant nodes of an NLR immune receptor network") OR TI=("Tempo and mode of morphological evolution are decoupled from latitude in birds") OR TI=("PhyloFisher: A phylogenomic package for resolving eukaryotic relationships") OR TI=("Global and national trends, gaps, and opportunities in documenting and monitoring species distributions") OR TI=("Perception of biological motion by jumping spiders") OR TI=("gen3sis: A general engine for eco-evolutionary simulations of the processes that shape Earth's biodiversity") OR TI=("Response thresholds alone cannot explain empirical patterns of division of labor in social insects") OR TI=("An oxytocin/vasopressin-related neuropeptide modulates social foraging behavior in the clonal raider ant") OR TI=("Population dynamics of Baltic herring since the Viking Age revealed by ancient DNA and genomics") OR TI=("Initial impressions of compatibility and mate value predict later dating and romantic interest") OR TI=("Development of mass allelic exchange, a technique to enable sexual genetics in Escherichia coli") OR TI=("Baboon perspectives on the ecology and behavior of early human ancestors") OR TI=("Delay effects on the stability of large ecosystems") OR TI=("Spatial turnover of soil viral populations and genotypes overlain by cohesive responses to moisture in grasslands") OR TI=("An insect brain organizes numbers on a left-to-right mental number line") OR TI=("Factors influencing terrestriality in primates of the Americas and Madagascar") OR TI=("Bending the curve: Simple but massive conservation action leads to landscape-scale recovery of amphibians") OR TI=("Fast acrobatic maneuvers enable arboreal spiders to hunt dangerous prey") OR TI=("The evolution of insular woodiness") OR TI=("Defensive symbiosis against giant viruses in amoebae") OR TI=("Accumulation and maintenance of information in evolution") OR TI=("Environmental context dependency in species interactions") OR TI=("Different classes of genomic inserts contribute to human antibody diversity") OR TI=("A single introduction of wild rabbits triggered the biological invasion of Australia") OR TI=("Signatures of adaptive evolution in platyrrhine primate genomes") OR TI=("Fine-scaled climate variation in equatorial Africa revealed by modern and fossil primate teeth") OR TI=("Wildlife susceptibility to infectious diseases at global scales") OR TI=("Sexual repurposing of juvenile aposematism in locusts") OR TI=("A set of principles and practical suggestions for equitable fieldwork in biology") OR TI=("Geographic patterns of koala retrovirus genetic diversity, endogenization, and subtype distributions") OR TI=("Lizards from warm and declining populations are born with extremely short telomeres") OR TI=("Concerted evolution of metabolic rate, economics of mating, ecology, and pace of life across seed beetles") OR TI=("The evolution of mating preferences for genetic attractiveness and quality in the presence of sensory bias") OR TI=("Glassfrogs conceal blood in their liver to maintain transparency") OR TI=("Rapid weed adaptation and range expansion in response to agriculture over the past two centuries") OR TI=("Madagascar's extraordinary biodiversity: Evolution, distribution, and use") OR TI=("Madagascar's extraordinary biodiversity: Threats and opportunities") OR TI=("Phosphoenolpyruvate reallocation links nitrogen fixation rates to root nodule energy state") OR TI=("The lower Cambrian lobopodian Cardiodictyon resolves the origin of euarthropod brains") OR TI=("Grazing and ecosystem service delivery in global drylands") OR TI=("Seventy years of tunas, billfishes, and sharks as sentinels of global ocean health") OR TI=("Hydraulic failure as a primary driver of xylem network evolution in early vascular plants") OR TI=("Refactored genetic codes enable bidirectional genetic isolation") OR TI=("Unprecedented fire activity above the Arctic Circle linked to rising temperatures") OR TI=("The control of carpel determinacy pathway leads to sex determination in cucurbits") OR TI=("PLANT SCIENCE Floral sex determination") OR TI=("Attenuated evolution of mammals through the Cenozoic") OR TI=("Evolution and antiviral activity of a human protein of retroviral origin") OR TI=("Disease outbreaks select for mate choice and coat color in wolves") OR TI=("Evolution of increased complexity and specificity at the dawn of form I Rubiscos") OR TI=("Sex-and age-dependent genetics of longevity in a heterogeneous mouse population") OR TI=("Molecular and cellular evolution of the primate dorsolateral prefrontal cortex") OR TI=("Genetic diversity loss in the Anthropocene") OR TI=("Termite sensitivity to temperature affects global wood decay rates") OR TI=("Exceptional preservation of organs in Devonian placoderms from the Gogo lagerstatte") OR

TI=("Codiversification of gut microbiota with humans") OR TI=("The genomic history and global expansion of domestic donkeys") OR TI=("Evolutionary gain and loss of a pathological immune response to parasitism") OR TI=("Symbiotic microbes aid host adaptation by metabolizing a deterrent host pine carbohydrate D-pinitol in a beetle-fungus invasive complex") OR TI=("Wild chimpanzee behavior suggests that a savanna- mosaic habitat did not support the emergence of hominin terrestrial bipedalism") OR TI=("Dental form and function in the early feeding diversification of dinosaurs") OR TI=("Coextinctions dominate future vertebrate losses from climate and land use change") OR TI=("Active DNA demethylation of developmental cis-regulatory regions predates vertebrate origins") OR TI=("TENT5 cytoplasmic noncanonical poly(A) polymerases regulate the innate immune response in animals") OR TI=("Chimera states among synchronous fireflies") OR TI=("Clustering of vomeronasal receptor genes is required for transcriptional stability but not for choice") OR TI=("A conserved module in the formation of moss midribs and seed plant axillary meristems") OR TI=("Genetic slippage after sex maintains diversity for parasite resistance in a natural host population") OR TI=("Flies trade off stability and performance via adaptive compensation to wing damage") OR TI=("The direct drivers of recent global anthropogenic biodiversity loss") OR TI=("Barriers to gene flow in the deepest ocean ecosystems: Evidence from global population genomics of a cosmopolitan amphipod") OR TI=("Diversity dependence is a ubiquitous phenomenon across Phanerozoic oceans") OR TI=("Megafauna extinctions produce idiosyncratic Anthropocene assemblages") OR TI=("1000 spider silkomes: Linking sequences to silk physical properties") OR TI=("Orientation pinwheels in primary visual cortex of a highly visual marsupial") OR TI=("Disease resistance in coral is mediated by distinct adaptive and plastic gene expression profiles") OR TI=("The genetic architecture of phenotypic diversity in the Betta fish (Betta splendens)") OR TI=("Floral color is not as simple as it once seemed") OR TI=("Ambush predation and the origin of euprimates") OR TI=("Proteotype coevolution and quantitative diversity across 11 mammalian species") OR TI=("Lost in translation: Molecular basis of reduced flower coloration in a self-pollinated monkeyflower (Mimulus) species") OR TI=("Temporal gene expression patterns in the coral Euphyllia paradivisa reveal the complexity of biological clocks in the cnidarian-algal symbiosis") OR TI=("Coevolution of the Ess1-CTD axis in polar fungi suggests a role for phase separation in cold tolerance") OR TI=("Survival fluctuation is linked to precipitation variation during staging in a migratory shorebird") OR TI=("Comparative environmental RNA and DNA metabarcoding analysis of river algae and arthropods for ecological surveys and water quality assessment") OR TI=("Quantifying thermal cues that initiate mass emigrations in juvenile white sharks") OR TI=("Mutagenesis alters sperm swimming velocity in Astyanax cave fish") OR TI=("A genotyping by sequencing approach can disclose Apis mellifera population genomic information contained in honey environmental DNA") OR TI=("Manatee calf call contour and acoustic structure varies by species and body size") OR TI=("Water provisioning increases caged worker bee lifespan and caged worker bees are living half as long as observed 50 years ago") OR TI=("Evolution of cross-tolerance in Drosophila melanogaster as a result of increased resistance to cold stress") OR TI=("Evidence for compositionality in baboons (Papio papio) through the test case of negation") OR TI=("Acoustic and visual cetacean surveys reveal year-round spatial and temporal distributions for multiple species in northern British Columbia, Canada") OR TI=("Living on the sea-coast: ranging and habitat distribution of Asiatic lions") OR TI=("The spread of Carpophilus truncatus is on the razor's edge between an outbreak and a pest invasion") OR TI=("A large-scale dataset reveals taxonomic and functional specificities of wild bee communities in urban habitats of Western Europe") OR TI=("Sperm tendency to agglutinate in motile bundles in relation to sperm competition and fertility duration in chickens") OR TI=("Factors influencing lion movements and habitat use in the western Serengeti ecosystem, Tanzania") OR TI=("A short exposure to a semi-natural habitat alleviates the honey bee hive microbial imbalance caused by agricultural stress") OR TI=("The dominant mesopredator and savanna formations shape the distribution of the rare northern tiger cat (Leopardus tigrinus) in the Amazon") OR TI=("Trophic provisioning and parental trade-offs lead to successful reproductive performance in corals after a bleaching event") OR TI=("Ornamental roses for conservation of leafcutter bee pollinators") OR TI=("Effects of vessel traffic and ocean noise on gray whale stress hormones") OR TI=("Investigation of the spermathecal morphology, reproductive strategy and fate of stored spermatozoa in three important thysanopteran species") OR TI=("Strategies of protected area use by Asian elephants in relation to motivational state and social affiliations") OR TI=("Influence of mating strategies on seminal material investment in crabs") OR TI=("Breeding and migration performance metrics highlight challenges for White-naped Cranes") OR TI=("The trophic niche of subterranean populations of Speleomantes italicus")
